# Quantifying Uncertainty and Robustness in a Biomathematical Model Based Patient-Specific Response Metric for Glioblastoma

**DOI:** 10.1101/325340

**Authors:** Andrea Hawkins-Daarud, Sandra K. Johnston, Kristin R. Swanson

**Affiliations:** Department of Neurosurgery, Mayo Clinic Arizona, Phoenix, AZ; Department of Radiology, University of Washington, Seattle WA

## Abstract

Glioblastomas, lethal primary brain tumors, are known for their heterogeneity and invasiveness. A growing literature has been developed demonstrating the clinical relevance of a biomathematical model, the Proliferation-Invasion (PI) model, of glioblastoma growth. Of interest here is the development of a treatment response metric, Days Gained (DG). This metric is based on individual tumor kinetics estimated through segmented volumes of hyperintense regions on T1-weighted gadolinium enhanced (T1Gd) and T2-weighted magnetic resonance images (MRIs). This metric was shown to be prognostic of time to progression. Further, it was shown to be more prognostic of outcome than standard response metrics. While promising, the original paper did not account for uncertainty in the calculation of the DG metric leaving the robustness of this cutoff in question. We harness the Bayesian framework to consider the impact of two sources of uncertainty: 1) image acquisition and 2) interobserver error in image segmentation. We first utilize synthetic data to characterize what non-error variants are influencing the final uncertainty in the DG metric. We then consider the original patient cohort to investigate clinical patterns of uncertainty and to determine how robust this metric is for predicting time to progression and overall survival. Our results indicate that the key clinical variants are the time between pre-treatment images and the underlying tumor growth kinetics, matching our observations in the clinical cohort. Finally, we demonstrated that for this cohort there was a continuous range of cutoffs between 94 and 105 for which the prediction of the time to progression and was over 80% reliable. While further validation must be done, this work represents a key step in ascertaining the clinical utility of this metric.

## 1 Introduction

Glioblastomas, lethal primary brain tumors, are known for their locally invasive nature where the tumor cells infiltrate far beyond the margins of the observed MRI abnormality [1]. At the time of this writing, clinicaltrials.gov lists 676 trials for interventional therapies of all phases for glioblastoma that have either been completed, suspended, or terminated and another 452 trials that are active in some form [2]. Despite this concerted effort, the standard-of-care for newly diagnosed glioblastoma patients has remained the same since 2005 [3].

A key reason for the apparent failure of these hundreds of clinical trials is the interpatient and intratumor heterogeneity of this disease. Ultimately, the success (or failure) of a clinical trial is defined by the difference in the median survival between the treated cohort and a current or historical control cohort rather than individual successes. This is because there is no known quantitative surrogate for assessing the impact of treatment in individual patients. MRI, the primary method of assessing the disease, is only able to show changes in the microenvironment due to the tumor (or therapy) rather than the tumor cells themselves. Pragmatically, this has resulted in no reliable generalization from MRI signals to tumor cell densities for use in determining therapeutic efficacy.

### Mathematical Models and Untreated Virtual Controls

A growing literature has begun demonstrating the clinical relevance of a patient-specific biomathematical model of glioblastoma growth [4, 5, 6, 7, 8]. This simple model, the Proliferation-Invasion (PI) model, can be written mathematically as:

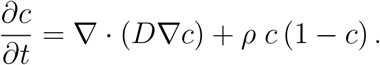

Here *c* is the tumor cell density, *D* is the net rate of invasion, and *ρ* is the net rate of proliferation. The output of this model is the spatial tumor cell density over time, which asymptotically approaches a traveling wave. The shape of the wave depends on the ratio of the two parameters, *D/ρ* while the speed of the wave depends on their product, 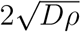. A number of groups have published methods to estimate these parameters [9, 10, 11]. Following [4], all methods abstract the abnormal regions on MRI to be representative of some tumor cell density threshold, 80% and 16% for the T1-weighted with gadolinium (T1Gd) and the T2-weighted sequences respectively. The Swanson group has demonstrated that patient-specific estimates of these parameters can be used in many contexts [6, 12, 13, 7, 8, 5]. Of interest to this paper, is their use in generating Untreated Virtual Controls (UVCs) for use in defining response metrics.

The response metric we focus on here is referred to as Days Gained (DG) and has been shown to be prognostic of both time to progression and overall survival when utilized at the first imaging time point after radiation therapy [14, 15]. It is calculated by 1) estimating the patient-specific growth kinetics through the PI model, 2) simulating the patients untreated tumor growth, 3) aligning the predicted growth with the observed pre-treatment sizes, 4) finding the time when the untreated tumor is predicted to be the size of the observed tumor after therapy and 5) subtracting it from the time corresponding to the post treatment observation. This final difference is the DG metric and represents how far back therapy pushed the tumor on its growth curve. This metric has been tested with individualized 3D anatomical simulations [14] and in simpler scenarios of spherical symmetry and linear growth [15]. In each case, it was observed that a cutoff could be found that statistically separated the patients into prognostically different“responders” vs. “non-responders.” However, to begin advocating for clinical use, more testing needs to be done both in terms of the generality and robustness of this metric. This paper addresses robustness, with particular focus on how uncertainty in the initial calculations of tumor size from MRI impacts the uncertainty in the DG metric.

## 2 Bayesian Methodology

To connect uncertainties in data with model predictions, we will utilize the Bayesian Framework. While the ultimate interest is the uncertainty in the DG metric, one must first quantify how the measurement error first propagates into uncertainty in the model parameters. This methodology relies on Bayes Theorem:

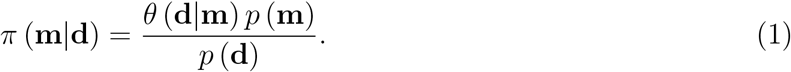

Here, **m** is interpreted as the model parameters and **d** as the observed data. The conditional probability π (**m|d**) is the posterior probability and can be thought of as the solution of a calibration problem. *p* (**m**) is the prior and represents the belief in the model parameters prior to observing the most recent data. The conditional probability *θ* (**d|m**) is the likelihood and captures what data one would expect to see if a particular set of model parameters were “truth”. *p* (**d**) captures the probability of observing the data given the specific modeling structure but is often just considered a normalizing constant.

We remark that Le et al. [11] also utilized a Bayesian formulation to estimate *D* and *ρ* with two critical differences. First, they considered the tumor in its anatomical location whereas we assume the tumor grows spherically-symmetrically. Second, Le et al. did not consider uncertainty in the original segmentation or their model’s initial condition. While the first difference means their individual model simulations are much more complicated, the second difference results in an uncertain “initial time point” of the first time point of imaging. This particular uncertainty greatly complicates the form of our likelihood.

Deterministic methods for calculating *D* and *ρ* rely on tumor segmentations from two MRI time points, with at least one time point including both a T1Gd and T2 sequence. For simplicity in this presentation, we will assume the first time point has both a T1Gd and T2 image and the second time point has only a T1Gd image. Then, for the current specific scenario of interest, the model parameters and data are defined as:

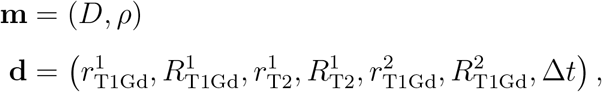

where the 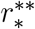’s and 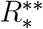’s represent the lower and upper bound of a measurement value, respectively, on the * MRI sequence and the ** day (first or second) and Δ*t* is the time between the first and second imaging dates.

### The Prior

We consider two different priors in this paper: 1) a uniform prior to allow maximal influence of the data, 2) a bivariate Gaussian prior fit to existing known parameter estimates from a large cohort of patients in the Swanson Lab Database. See supplementary material.

### The Likelihood

To define the likelihood, we first assume that the T1Gd and T2 MRIs respectively reflect the 80% and 16% cell density thresholds. We then consider two sources of uncertainty: 1) The ability of the MRI to accurately recapitulate the truth, 2) the ability of a human for measuring the truth that the MRI is showing. The first uncertainty results from various issues such as images being composed of two-dimensional slices with non-zero thickness and that MRI machines will use different acquisition parameters. While it is difficult to define these uncertainties with rigor, the framework presented here is easily modifiable to account for any additional information. Regarding the second type of uncertainty, we have a large number of images that have been measured twice allowing for some approximation of this variability.

Due to the uncertainty in when the first time point of observation is relative to the model’s output, our likelihood is substantially more complicated than a traditional Gaussian. It is composed of multiple bivariate probability distribution functions of radius, *r*, and time, *t*, and takes the final form:

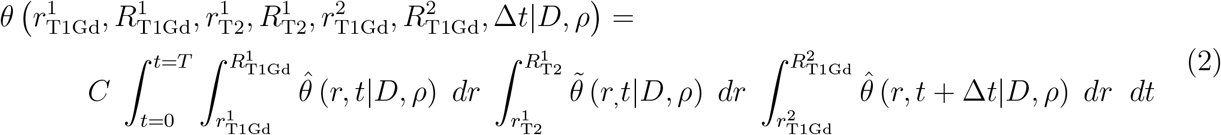

In this equation, *C* is a normalization constant that must be calculated for each *D* and *ρ* and the functions 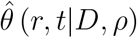 and 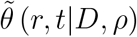 capture the probability of the T1Gd or T2 MRI, respectively, showing an imaging abnormality of size *r* at simulation time *t* for a given *D* and *ρ*. For each simulation time *t*, we assume this probability is a triangle distribution around the model predicted radius. See supplemental material.

In words, the output of this function is the sum of the probability from each possible initial time point of observing the range of possible radial measurements all at once.

### The Posterior

The posterior is considered the solution to the calibration problem and can be written down as being proportional to the product of the prior and likelihood.

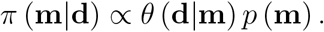

While the functional form of the posterior is easy to write down, the complicated likelihood function, which must be computed to evaluate the posterior at any given point, makes analytical assessment impossible.

### Uncertainty Propagation to DG Response Metric

To propagate the uncertainty from the calibration to an individual’s DG response metric, we must incorporate additional uncertainty coming from the measurement of the post-therapy image. See Figure 1.

**Figure 1:**
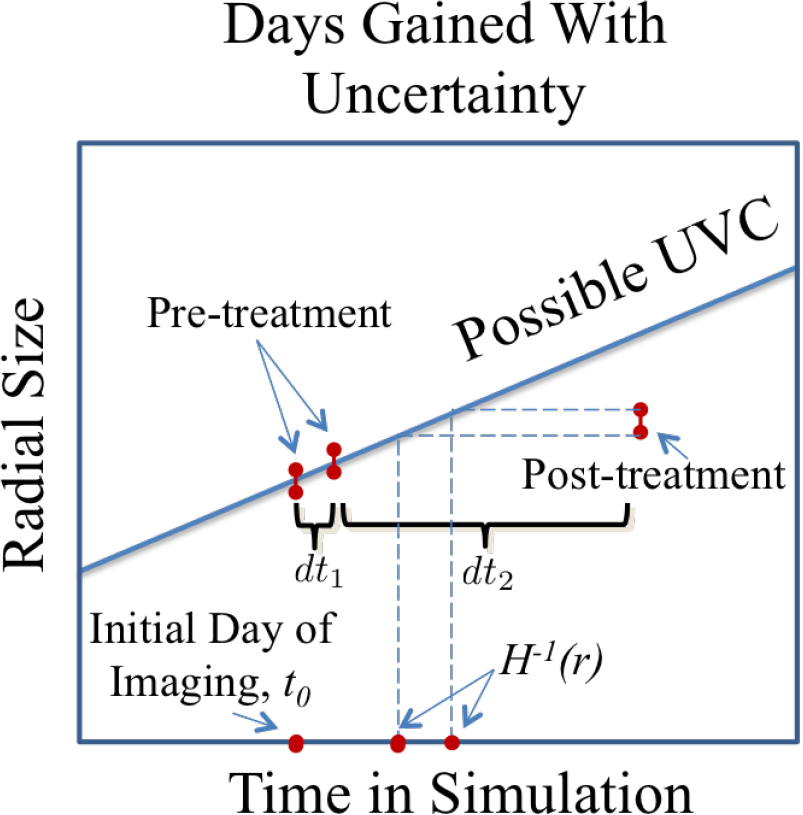
Illustration of components required to calculate the probability of the DG metric.

Thus, the final probability distribution we must calculate is: 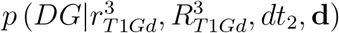. This is evaluated utilizing the total law of probability with the previously calculated posterior and the probability of the initial time point. Code was written in MATLAB [16]. See the supplement for additional numerical details.

## 3 Methods

### 3.1 Identifying Non-Error Influences on Uncertainty in DG

There are numerous variabilities between any two given patients that could influence the ultimate uncertainty in their DG metric. We consider three such factors: 1) the individual growth kinetics, 2) the size of tumor when therapy began, and 3) the length of time between the two pre-treatment imaging time points.

#### Different Growth Kinetics

To understand how these factors may all interact together we ran 4 simulations with different growth kinetics from which we generated synthetic data representative of different situations related to 1) and 2). Specifically, we ran simulations with the (*D* [mm^2^/yr],*ρ* [1/yr]) pairs of (3, 3), (30, 30), (30, 3), and (3, 30). Note, the first two pairs have the same “slope” of their traveling wave while the last two have the same velocity.

#### Different Size at Therapy Initiation

For each (*D*, *ρ*) pair, we made the assumption that a standard course of radiation therapy lasting for six weeks was initiated when the tumor reached different radial sizes, 1 cm, 2 cm, and 3 cm. Then, to minimize variables influencing the uncertainty, we assumed that 50 days after the last pre-treatment image time point, all tumors were observed at a size of 1.5 cm.

#### Different

Δ*t* These 12 scenarios, four (*D*, *ρ*) pairs with three different sizes at treatment initiation, were used to generate synthetic “observed” data corresponding to different time periods between the pre-treatment time points. We identified the size just prior to the start of therapy and then either 10, 25, or 40 days prior to this simulation date as the first pre-treatment date.

The Bayesian methodology described previously was applied to each of these sets of generated data, both with a uniform and Gaussian prior, assuming an interobserver variability of 0.3 mm in measured radius.

### 3.2 Interpreting DG within a cohort of patients under uncertainty

Our Bayesian algorithm was implemented for each patient in the original cohort from [15] with both priors to generate individual predictions of the probability of the patient’s DG score. Images where two measurements were available provided upper and lower bounds for the measurement uncertainty. Images with only one measurement were assigned 0.3 mm radius uncertainty.

#### Patients

The 63 patients analyzed in the previously published paper [15] were again analyzed in this paper for direct comparison. Criteria for inclusion are 1) over 18, 2) GBM diagnosis, 3) two pre-treatment imaging timepoints with a minimum of 4 days in between, and 4) upfront therapy inclusive of radiation therapy. Survival and time to progression were updated as available.

#### Quantifying Robustness

1000 samples were generated from each patients’ DG probability and collated to generate 1000 realizations of this entire cohort’s DG values. For each cohort realization, we iterated through all possible cutoffs to define responders and evaluated the log-rank of the corresponding Kaplain-Meyer curves corresponding to both the time to progression and overall survival, storing the p-values. We then determined how many times each cutoff resulted in a significant outcome difference at the p=0.05 level. High counts are correlated with robustness of the cutoff to uncertainty. Results with both priors are considered.

## 4 Results

### Size of tumor at therapy initiation seems to have no impact on uncertainty

In considering the four (*D*, *ρ*) pairs and all simulated sizes of tumor at the start of radiation, the resulting posterior and DG probability density function were independent of the size of the tumor when therapy began. Results not shown.

### Uncertainty decreases in posterior for larger

Δ*t*. Figure 2 shows the posterior using a uniform prior and the DG probability density function for all four (*D*, *ρ*) pairs, with treatment starting when the tumor was 2 cm in size, but with Δ*t* = 10 or 40 days. Red dots indicate the true values. In each case, the posterior uncertainty decreases as you have more time between images. We note that for small Δ*t*, the uncertainty demonstrated in the posterior can encompass almost all possible velocities.

**Figure 2:**
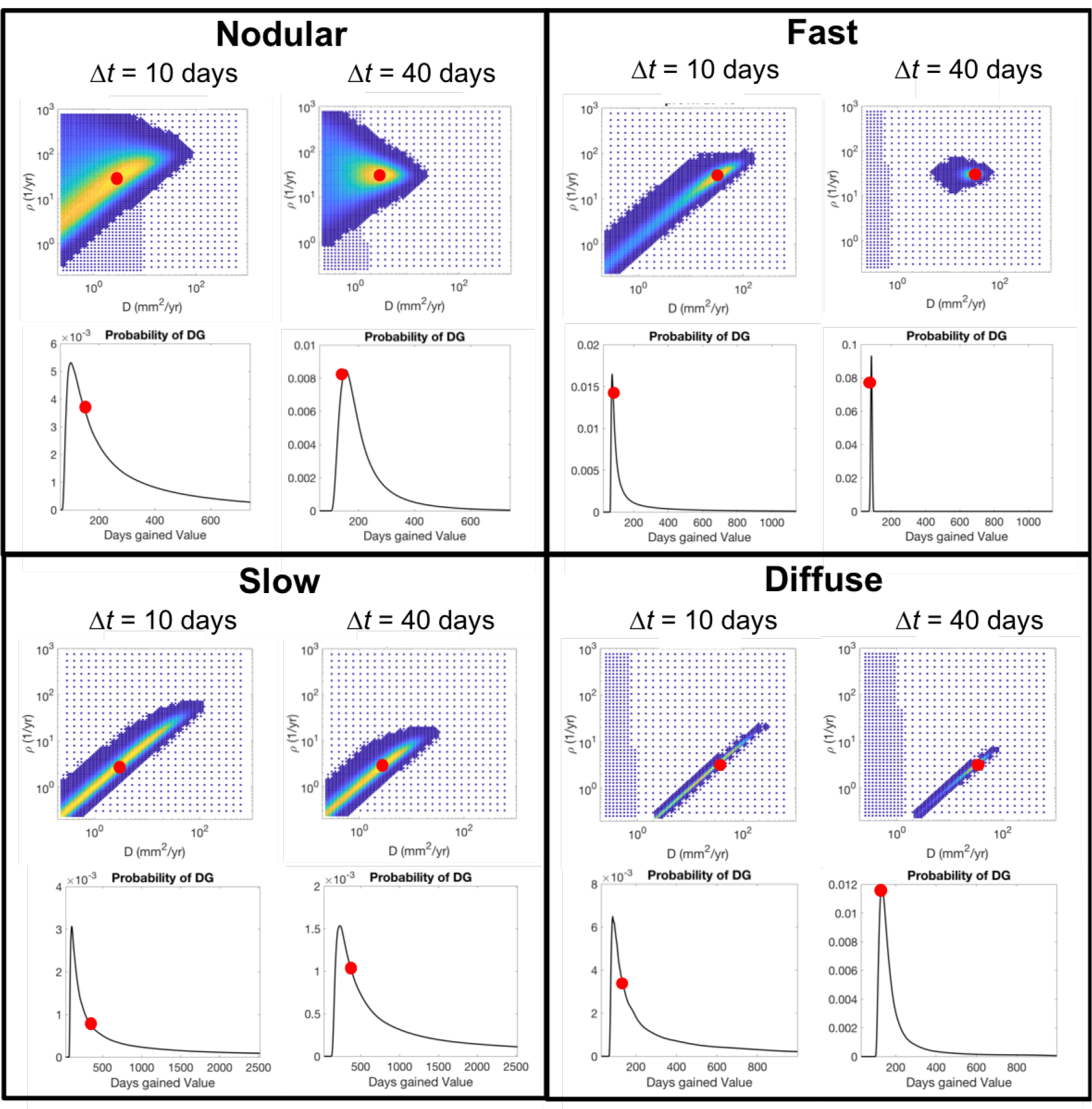
Synthetic Examples: Uncertainty in the posterior probability density function uniformly decreases as the time between the pre-treatment images increases. Shorter time intervals result in a large degree of uncertainty in the posterior in the velocity of the traveling wave. We also see that the underlying tumor growth kinetics influence the degree to which the time between the images matters. This has an interesting impact on the resultant probability of DG which is dependent on the underlying tumor growth kinetics. Specifically the slower growing tumors, with a higher degree of uncertainty in their posterior in both scenarios, seem to have a wider distribution, or greater uncertainty, for larger Δ*t*, but the probability of the truth increases as the distribution widens. This is due to a bias towards the DG values predicted by the faster tumors given equal weight in the posterior.

### Degree of uncertainty does depend on underlying tumor growth kinetics

Figure 2 also demonstrates that the degree of uncertainty does depend on the underlying tumor growth kinetics. Specifically, slower tumors have a larger uncertainty in velocity, no matter the Δ*t*. Additionally, the nodularity, or steepness of the traveling wave, influences the uncertainty as the nodular example exhibits a wider distribution.

### Apparent uncertainty in DG dependent on underlying tumor growth kinetics

In Figure 2, one can see that while the probability of the true DG value increases as Δ*t* increases, the width of the distribution sometimes increases. This is because higher velocity growth patterns result in a tighter prediction of DG values. Thus, when a broad range of fast and slow tumors are represented equally in the posterior, the DG will be biased towards the DG values associated with the faster growth. This bias diminishes as uncertainty in the posterior is reduced.

### Posteriors can be greatly influenced by the Gaussian Prior

When the uncertainty is high in the likelihood, the Gaussian prior will have a significant impact on the posterior, as seen in Figure 3. While it may seem to reduce the uncertainty, it may also skew the probability away from the true values. A more certain likelihood is less influenced by the prior.

**Figure 3:**
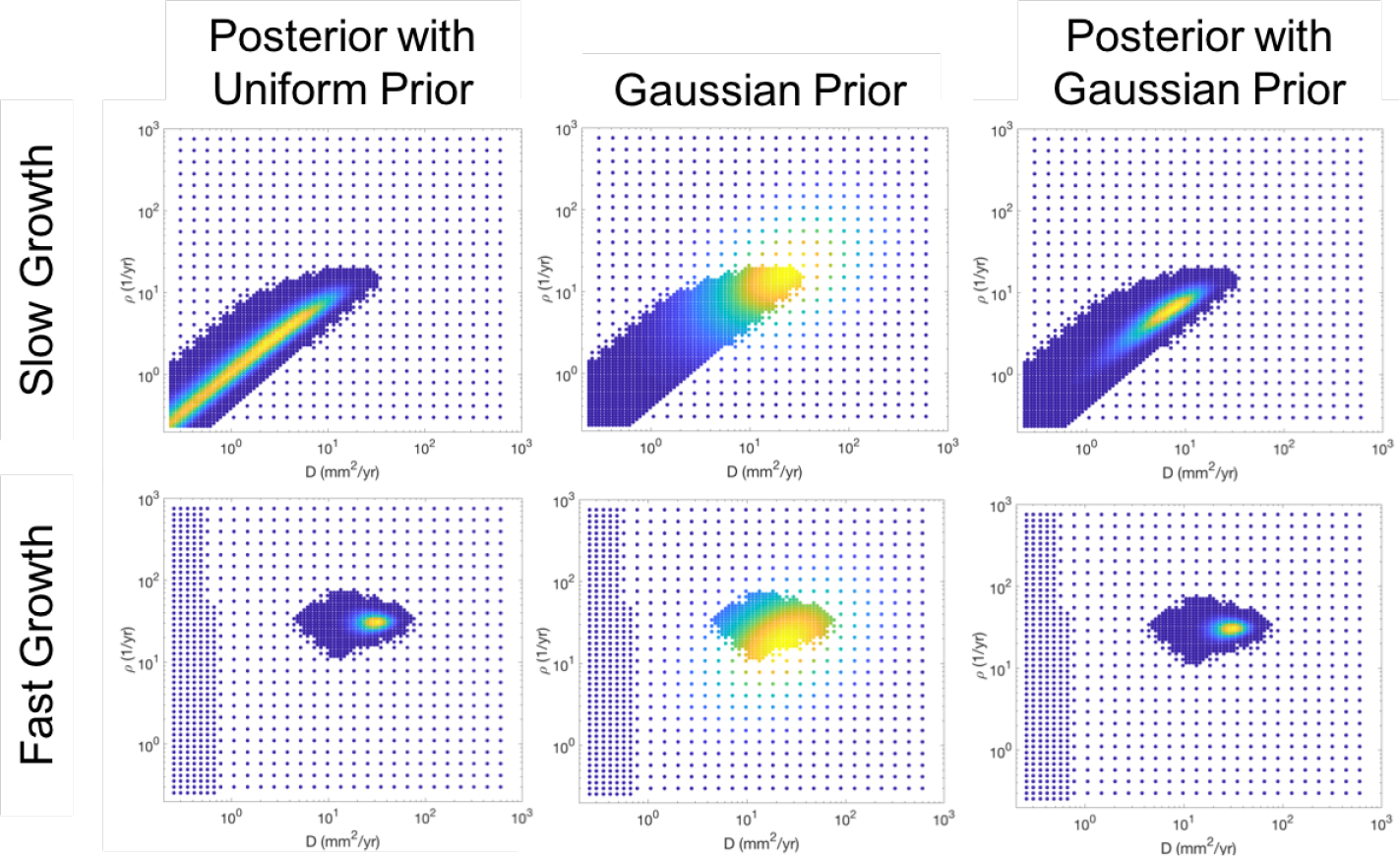
Influence of the Prior. The top and bottom rows correspond to the slow and fast growing tumor respectively, each with 40 days between the pre-treatment images. The left column shows the posterior when a uniform prior is assumed and is thus equivalent to the likelihood, this is the same as is seen in Figure 2. The middle column shows the value of the Gaussian prior at the same node points and the right column shows the posterior when the Gaussian prior is utilized. When the uncertainty is high in the likelihood, the Gaussian prior can significantly reduce the uncertainty, but also biases it towards where the prior has high probability. When there is less uncertainty in the prior, such as for the fast growing tumor, the prior has less influence.

### Uncertainty in DG metric as realized in patient cohort

Figure 4 shows a summary of the results from running the Bayesian methodology on the original cohort of patients presented in [15] with both a uniform and Gaussian prior. Each dot represents a patient with the (x,y) coordinate pair indicating the maximum likely *D* and *ρ* for that patient under this framework. The relative width of the 90% confidence interval of the DG probability distribution is indicated by the size of the dot on Figure 4. Patients were further identified based on their Δ*t*, indicated by color. As we expect given Figure 2, the larger, more uncertain, dots are in the lower left corner irrespective of color. It is also evident that while the uncertainty is decreased in general from the left to right plot, the dots also move significantly as they are pulled towards the high probability regions of the Gaussian prior.

**Figure 4:**
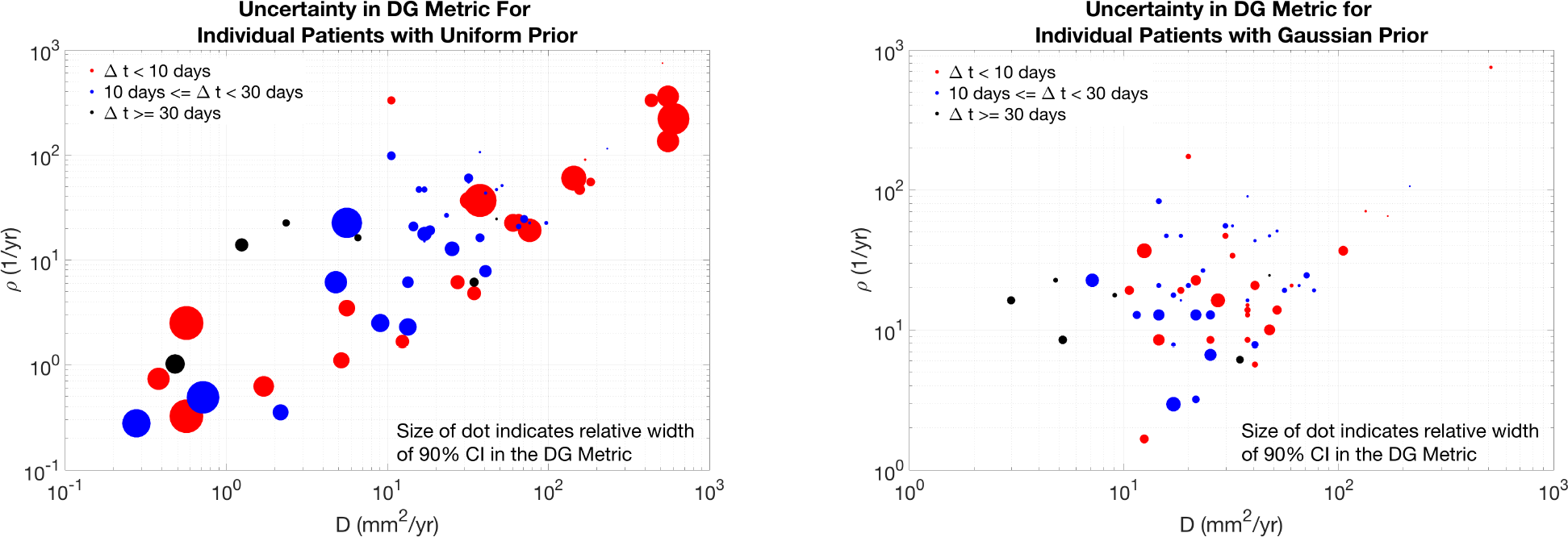
Illustration of Uncertainty in DG value for original cohort studied in [15] for both uniform and Gaussian priors. The (x,y) coordinate pair of each circle indicates the maximum likely *D* and *ρ* for that patient under the Bayesian framework with the indicated prior. The size of the dot indicates the relative width of the 90% confidence interval of the DG probability density function, while the color indicates the length of time between the pre-treatment images. As expected from Figure 2, the slower growing tumors, those in the bottom left corner, do tend to have the larger uncertainty in spite of the time between the pre-treatment images. This also shows how the Gaussian prior biases the *D* and *ρ* selection towards the middle of the parameter space and reduces uncertainty in the DG metric.

### Quantify Robustness

The insets in Figure 5 illustrate the logrank p-value for each cutoff within each cohort realization, p-value of 1 is yellow and 0 is dark blue for the four scenarios of predicting time to progression and overall survival utilizing either the uniform the Gaussian prior. The histogram shows the number of times the specific DG cutoff resulted in a statistical difference in outcome between the two cohorts. In general, the DG metric was better able to serve as a prognostic indicator for the time to progression as the counts were higher. In particular, a similar range of DG values for either the uniform (92 to 105) or the Gaussian prior (94 to 106) was seen to be prognostic more than 80% of the time.

**Figure 5:**
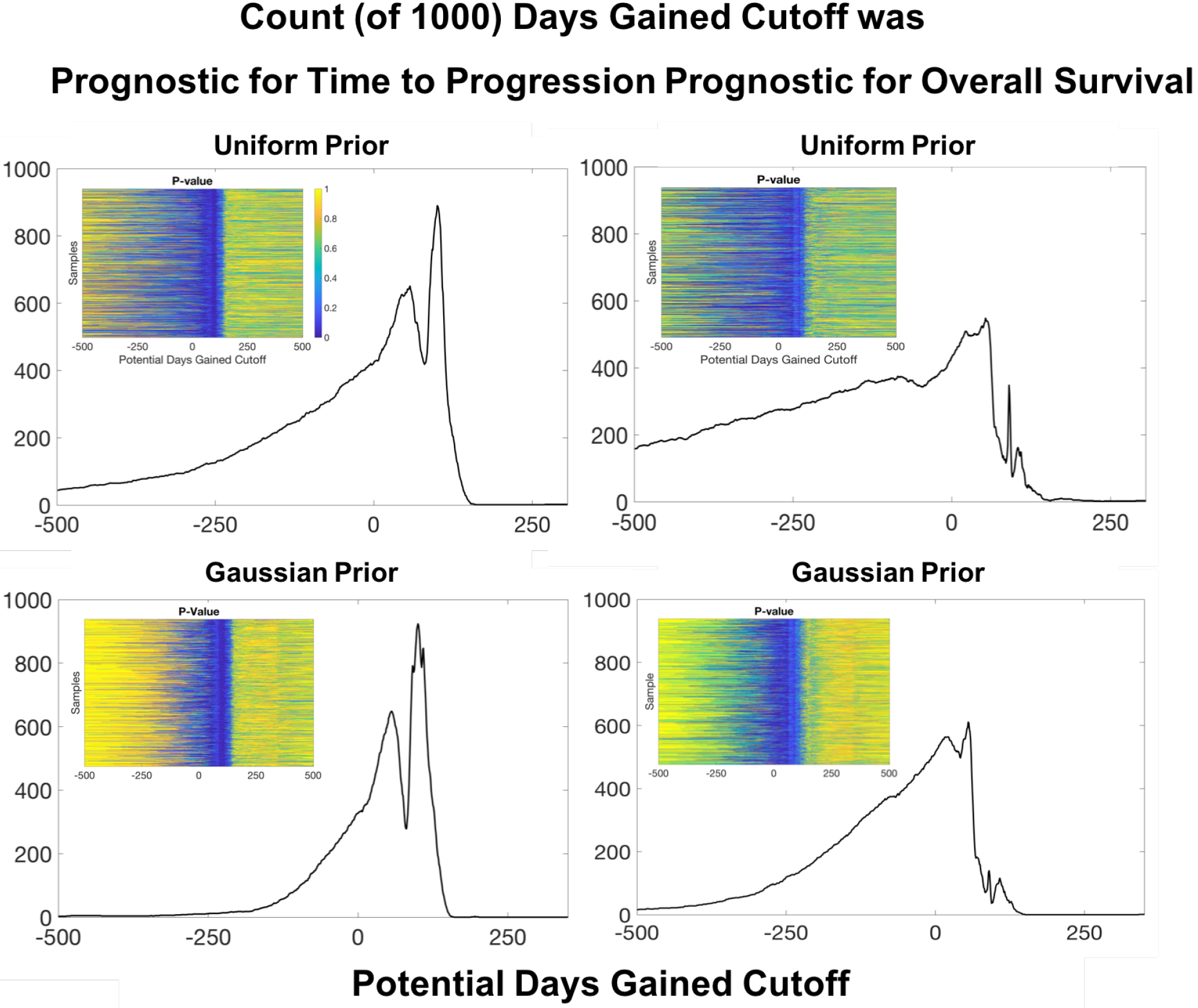
Quantification of Robustness: 1000 cohort samples were generated of Days Gained values from the individual patient distributions resulting from either a uniform or Gaussian prior. For each sample cohort we assessed at each possible cutoff the p-value associated with a log-rank test of differences in both time to progression and overall survival. By summing the number of times each cutoff was found to be statistically significant at the p=0.05 level, we demonstrated there is a good range of values for which the Days Gained cutoff is 80% of the time prognostic for time to progression, 93 to 105 for the uniform prior and 94 to 106 for the Gaussian prior. Doing the same analysis, we saw there was no cutoff that provided the same robustness in prognosis for overall survival, though there are some cutoffs that are prognostic about half the time.

## 5 Discussion and Conclusion

Here we have implemented a Bayesian framework to assess the uncertainty in a promising biomathematical model based response metric, Days Gained. We have ignored model uncertainty and focused solely on uncertainty from measurements of MRI abnormalities. By using synthetic data and considering both a uniform and Gaussian prior, we were able to characterize non-error influencing factors on the uncertainty. Using the patients in the original cohort presented in [15], we verified these patterns of uncertainty manifested in a clinical setting.

The strength of the Bayesian framework in this context is that it allows one to consider how the uncertainty, both the “from where” and “how much”, ultimately impacts a prediction and thus its robustness. The often criticized weakness of Bayesian methods is the choice of the prior, as it has significant potential to influence results (See Figure 3). In situations such as those presented here, where there is not a large amount of data for any given patient, the use of an informative prior reduces the uncertainty but is not guaranteed to increase the accuracy. As in all cases, confidence in the prior comes from confidence in how well the current sample is represented by the samples used to define the prior. The Gaussian prior used here, is defined using a large, heterogeneous population of GBM patients with previously estimated tumor growth kinetics. While it is encouraging and notable that the test for robustness (Figure 5 ultimately yielded the same range of prognostic cutoffs for both priors, we believe future efforts at refining the prior based on clinical features will enhance these results.

This work represents a critical step in translating mathematical models into clinical settings. While the obvious next step is to run this same investigation with a larger cohort, these preliminary results using the DG metric suggest that clinical trials may be able to use a surrogate for time to progression in determining therapeutic success. If true, this would revolutionize research in glioblastoma as clinical trials would be able to open and close more quickly. Zooming in, it could also revolutionize the treatment paradigm for individual patients by letting them make more informed decisions with their neurooncologists. Uncertainty is always present. But if the sensitivity of critical predictions can be quantified, the fear in making decisions can be minimized.

## Acknowledgements

The authors gratefully acknowledge the funding support of this work through the NSF 1122322, the NIH (R01CA16437, R01NS060752,U54CA210180, U54CA143970, U54193489, U01CA220378), the James S. McDonnell Foundation and the Ben & Catherine Ivy Foundation.

## References

[1] D. L. Silbergeld and M. R. Chicoine, “Isolation and characterization of human malignant glioma cells from histologically normal brain,” J. Neurosurgery, vol. 86, pp. 525–531, 1997.

[2] “Clinicaltrials.gov.” Accessed: 2018-04-26, Search Term “Glioblastoma”.

[3] R. Stupp, W. P. Mason, M. J. van den Bent, M. Weller, B. Fisher, M. J. Taphoorn, K. Belanger, A. A. Brandes, C. Marosi, U. Bogdahn, J. Curschmann, R. C. Janzer, S. K. Ludwin, T. Gorlia, A. Allgeier, D. Lacombe, J. G. Cairncross, E. Eisenhauer, and R. O. Mirimanoff, “Radiotherapy plus concomitant and adjuvant temozolomide for glioblastoma,” New England Journal of Medicine, vol. 352, no. 10, pp. 987–996, 2005. PMID: 15758009.

[4] K. R. Swanson, R. C. Rostomily, and E. C. Alvord Jr, “A mathematical modelling tool for predicting survival of individual patients following resection of glioblastoma: a proof of principle,” British Journal of Cancer, vol. 98, p. 113119, 2008.

[5] R. Rockne, J. K. Rockhill, M. M. a/nd A M Spence, I. Kalet, K. Hendrickson, A. Lai, T. Cloughesy, E. C. A. Jr, and K. R. Swanson, “Predicting the efficacy of radiotherapy in individual glioblastoma patients in vivo: a mathematical modeling approach,” Physics in Medicine & Biology, vol. 55, no. 12, p. 3271, 2010.

[6] A. L. Baldock, S. Ahn, R. C. Rockne, S. Johnston, M. L. Neal, D. Corwin, K. Clark-Swanson, G. Sterin, A. D. Trister, H. Malone, V. Ebiana, A. M. Sonabend, M. M. Mrugala, J. K. Rockhill, D. L. Silbergeld, A. Lai, T. F. Cloughesy, G. M. McKhann II, J. N. Bruce, R. C. Rostomily, P. Canoll, and K. R. Swanson, “Patient-specific metrics of invasiveness reveal significant prognostic benefit of resection in a predictable subset of gliomas,” PLoS ONE, vol. 9, no. 10, p. e99057, 2014.

[7] P. R. Jackson, J. Juliano, A. Hawkins-Daarud, R. C. Rockne, and K. R. Swanson, “Patient-specific mathematical neuro-oncology: Using a simple proliferation and invasion tumor model to inform clinical practice,” Bulletin of Mathematical Biology, vol. 77, pp. 846–856, May 2015.

[8] C. A. Rayfield, F. Grady, G. D. Leon, R. Rockne, E. Carrasco, P. Jackson, M. Vora, S. K. Johnston, A. Hawkins-Daarud, K. R. Clark-Swanson, S. Whitmire, M. E. Gamez, A. Porter, L. Hu, L. Gonzalez-Cuyar, B. Bendok, S. Vora, and K. R. Swanson, “Distinct phenotypic clusters of glioblastoma growth and response kinetics predict survival,” JCO Clinical Cancer Informatics, no. 2, pp. 1–14, 2018.

[9] K. R. Swanson, E. C. Alvord Jr, J. D. Murray, and R. C. Rockne, “Method and system for characterizing tumors,” 2013. U.S. Patent US8571844B2.

[10] A. Amelot, C. Deroulers, M. Badoual, M. Polivka, H. Adle-Biassette, E. Houdart, A. F. Carpentier, S. Froelich, and E. Mandonnet, “Surgical decision making from image-based biophysical modeling of glioblastoma: Not ready for primetime,” Neurosurgery, vol. 80, no. 5, pp. 793–799, 2017.

[11] M. Lê, H. Delingette, J. Kalpathy-Cramer, E. R. Gerstner, T. Batchelor, J. Unkelbach, and N. Ayache, “MRI Based Bayesian Personalization of a Tumor Growth Model,” IEEE Transactions on Medical Imaging, vol. 35, pp. 2329–2339, Apr. 2016.

[12] A. L. Baldock, K. Yagle, D. E. Born, S. Ahn, A. D. Trister, M. Neal, S. K. Johnston, C. A. Bridge, D. Basanta, J. Scott, H. Malone, A. M. Sonabend, P. Canoll, M. M. Mrugala, J. K. Rockhill, R. C. Rockne, and K. R. Swanson, “Invasion and proliferation kinetics in enhancing gliomas predict idh1 mutation status,” Neuro-Oncology, vol. 16, no. 6, pp. 779–786, 2014.

[13] A. L. Baldock, R. C. Rockne, A. D. Boone, A. Neal, Maxwell L. Hawkins-Daarud, D. M. Corwin, C. A. Bridge, L. A. Guyman, A. D. Trister, M. M. Mrugala, J. K. Rockhill, and K. R. Swanson, “From patient-specific mathematical neuro-oncology to precision medicine,” Frontiers in Oncology, vol. 3, no. 62, 2013.

[14] M. L. Neal, A. D. Trister, T. Cloke, R. Sodt, S. Ahn, A. L. Baldock, C. A. Bridge, A. Lai, T. F. Cloughesy, M. M. Mrugala, J. K. Rockhill, R. C. Rockne, and K. R. Swanson, “Discriminating survival outcomes in patients with glioblastoma using a simulation-based, patient-specific response metric,” PLoS ONE, vol. 8, no. 1, p. e51951, 2013.

[15] M. L. Neal, A. D. Trister, S. Ahn, A. L. Baldock, C. A. Bridge, L. Guyman, J. Lange, R. Sodt, T. Cloke, A. Lai, T. F. Cloughesy, M. M. Mrugala, J. K. Rockhill, R. C. Rockne, and K. R. Swanson, “Response classification based on a minimal model of glioblastoma growth is prognostic for clinical outcomes and distinguishes progression from pseudoprogression,” Cancer Research, vol. 73, no. 10, pp. 2976–2986, 2013.

[16] MATLAB, *version 7.10.0 (R2017b)*. Natick, Massachusetts: The MathWorks Inc., 2017.

